# Season-long infection of diverse hosts by the entomopathogenic fungus *Batkoa major*

**DOI:** 10.1101/2021.12.14.472666

**Authors:** Andrii P. Gryganskyi, Jacob Golan, Ann E. Hajek

## Abstract

Populations of the entomopathogenic fungus *Batkoa major* were analyzed using sequences of four genomic regions and evaluated in relation to their genetic diversity, insect hosts and collection site. This entomophthoralean pathogen killed numerous insect species from 23 families and five orders in two remote locations during 2019. The host list of this biotrophic pathogen contains flies, true bugs, butterflies and moths, beetles, and barkflies. Among the infected bugs (Order Hemiptera), the spotted lanternfly (*Lycorma delicatula*) is a new invasive planthopper pest of various woody plants that was introduced to the USA from Eastern Asia. A high degree of clonality occurred in the studied populations and high gene flow was revealed using four molecular loci for the analysis of population structure. We did not detect any segregation in the population regarding host affiliation (by family or order), or collection site. This is the first description of population structure of a biotrophic fungus-generalist in the entomopathogenic Order Entomophthorales. This analysis aimed to better understand the potential populations of entomopathogen-generalists infecting emerging invasive hosts in new ecosystems.

## Introduction

Species in the Entomophthorales are predominantly arthropod pathogens, serving important ecological roles ranging from modifying host behavior to regulating population dynamics (1–4). However, host range among entomophthoralean species is poorly understood, complicated by limited information on species identities of both fungal pathogens and arthropod hosts. Moreover, advances in sequencing technologies have revealed the presence of several species complexes, for example resulting in what was once considered a species with multiple hosts in fact being several cryptic species with distinctive host specificities (e.g., the *Entomophaga maimaiga* species complex (5), and the *Entomophthora muscae* species complex (6)). Additional complications arise when entomophthoralean-arthropod species combinations tested in the lab demonstrate pathogenicity, although field studies often reveal a narrower host range (i.e., ecological host range) than the lab host range (physiological host range (7)). Therefore, to understand the dynamics of diseases caused by entomophthoralean fungi in arthropod populations, it is critically important to identify the spectrum of potential arthropod hosts.

*Batkoa* is an excellent example of an entomophthoralean genus whose recent phylogenetic revision (8,9) allows for careful comparisons with arthropod hosts. The genus was first described in 1989 (10) and now includes ten species (11). Although at one point divided across six genera, recent phylogenetic studies provide parallel evidence that *Batkoa* is a single and distinct genus (9,12). Across all *Batkoa* species, the host range of *B. major* is among the best documented, with several reported host associations; in a worldwide compendium of entomophthoralean species, Bałazy (2) lists *B. major* as occurring in North and South America, as well as in Europe and Asia, and that it was “infecting several insect species of different orders,” including a ptilodactylid (Coleoptera), tipulids (Diptera), aphids (Hemiptera), and an ichneumonid (Hymenoptera). In 2018, *B. major* was also found alongside *Beauveria bassiana* (another entomopathogenic fungal species), co-infecting populations of the invasive spotted lanternfly (*Lycorma delicatula*, Fulgoridae, Hemiptera (13). This invasive fulgorid planthopper is only distantly related to native insects in the area of the co-epizootic (i.e., there are no native species in the same family, the Fulgoridae, in this area). At the time of the 2018 epizootics, *B. major* had only been cited from North America one time since its description in 1888 (8).

This study began with the goal of identifying the native reservoir hosts for *B. major*, a poorly known pathogen causing epizootics in outbreak populations of a new invasive. Based on trends in host range in the Entomophthorales, it was assumed that there would be few native host species and that these would predominantly belong to the order Hemiptera. We present results of a survey of naturally occurring infections in northeastern US forests that was conducted to identify hosts of *B. major*. We hypothesized that *B. major* in sampled locations is genetically diverse, but that it is not a species complex and that its populations are mostly clonal. Subsequent analyses investigated the genetic diversity and population structure of *B. major* to evaluate the potential for host specific clones and gene flow among collection sites.

## Materials and methods

### Sample collection and fungal isolation

Native insect populations were sampled in an Alleghany mixed hardwood forest near Ithaca, New York. On nine days between 19 June and 14 September 2019, cadavers of insects killed by entomophthoralean fungi were collected in Danby State Forest, Tompkins County, New York. Collections were made along the Abbott Loop hiking trail between 42.315636, -76.495048 / 42°18’56.3”N 76°29’42.2”W and 42.295817, -76.486345 / 42°17’44.9”N 76°29’10.8”W. Native insects killed by entomophthoralean fungi were also collected along the borders of the Angora Fruit Farm (40°21’30.6”N, 75°53’00.4”W), Berks County Parks and Recreation, Pennsylvania on 19 September 2019, near *Ailanthus altissima* (tree of heaven; the preferred host tree for *L. delicatula*) and in the adjacent hardwoods. At both sites, all sides of leaves, twigs, and branches from the ground to 2.5 m were carefully surveyed for dead insects. Arthropod cadavers were placed in 29 ml clear plastic cups containing 5 ml of 1.5% water agar and were transported to the laboratory at 4°C. Collected insects were from low density populations of native species. Collection trips were made within 24-48 hours after rainfall and collections took place over a period of two to three hours per site. The two sample sites are approximately 220 km from each other. In 2019, we collected a total of 213 insects that appeared to have been killed by entomophthoralean fungi. Most entomophthoralean species are notoriously difficult to isolate so collections resulted in a total of 67 samples of *B. major* that could be used for molecular analysis.

In the laboratory cadavers that were sporulating or were ready to sporulate were moved to high humidity enclosures at room temperature. Each cadaver was separately covered with the base of a 60 mm petri dish containing malt extract agar (MEA; 30 g malt extract, 20 g agar, 1 L distilled water) to allow “ascending” conidia to be collected on the MEA (14). After approximately 6 hours, petri dishes with conidia were removed. Cadavers that had not yet sporulated were left under high humidity at 20°C overnight and were irregularly checked for sporulation for a total of 48 h. After conidia had been collected or at 48 h, the body of each insect was stored at -20°C and subsequently examined to morphologically identify arthropod host species.

All MEA plates with conidia were maintained at 20°C. Once conidia had begun to germinate, thin sections of MEA containing hyphae were excised and placed in 35 mm petri dishes containing 1.5 ml 95% Grace’s insect medium (Lonza, Walkersville, MD) and 5% fetal bovine serum (Life Technologies, Grand Island, NY). Once hyphal growth was evident, hyphae were transferred to egg yolk Sabouraud maltose agar (EYSMA (14)) in 100 mm petri dishes. When cultures were mature, they were frozen in 10% glycerol in 2 ml cryotubes at -80°C, using a CoolCell Freezing System (Corning, NY) and deposited in the ARSEF culture collection (Table 1).

**Table 1..** Insect hosts collected in Pennsylvania and New York State, infected by *Batkoa major* in 2019-2020.

| Collection |  | Host |  |  | ARSEF #<br>(HLBio#) | GenBank accession # |  |  |  |
| --- | --- | --- | --- | --- | --- | --- | --- | --- | --- |
| Site | Date | Order | Family or Suborder | Species |  | ITS1 | ITS2 | 28S | RPB2 |
| AFF | 9/10 2018 | Hemiptera | Fulgoridae | *Lycorma delicatula* | Bat13769 | OL335101 | OL335159 | OL332699 | OL624704 |
| DSF | 6/27/2019 | Diptera | Rhagionidae | * | 14421 (Bat42) | OL335043 | OL335102 | OL332659 | OL624643 |
| DSF | 6/27/2019 | Hemiptera | Cixiidae | *Cixius* sp. | 14420 (Bat73) | OL335044 | OL335103 | OL332659 | OL624644 |
| DSF | 6/27/2019 | Coleoptera | Elateridae | *Athous brightwelli* | 14426 (Bat83) | OL335045 | OL335104 | OL332659 | OL624645 |
| DSF | 6/27/2019 | Coleoptera | Cantharidae | *Rhagonycha fraxini* | (Bat86) | OL335046 | OL335105 | OL332637 | OL624646 |
| DSF | 6/27/2019 | Diptera | Dolichopodidae | *Gymnopterus* sp. | (Bat88) | OL335047 | OL335106 | OL332638 | OL624647 |
| DSF | 6/27/2019 | Coleoptera | Elateridae | *Athous brightwelli* | 14444 (Bat90) | OL335048 | OL335107 | OL332639 | OL624648 |
| DSF | 6/27/2019 | Coleoptera | Tenebrionidae | *Isomira sericea* | (Bat91) | OL335049 | - | OL332640 | OL624649 |
| DSF | 7/10/2019 | Diptera | Lauxaniidae | *Homoneura inserta* | (Bat120) | OL335050 | OL335108 | OL332641 | OL624650 |
| DSF | 7/10/2019 | Diptera | Sciaridae | * | (Bat121) | - | - | OL332642 | - |
| DSF | 7/10/2019 | Diptera | Rhagionidae | *Symphoromyia* sp. | (Bat123) | OL335051 | OL335109 | OL332643 | OL624651 |
| DSF | 7/10/2019 | Diptera | Rhagionidae | * | (Bat124) | OL335052 | OL335110 | OL332644 | OL624652 |
| DSF | 7/10/2019 | Diptera | Rhagionidae | * | 14427 (Bat132) | OL335053 | OL335111 | OL332645 | OL624653 |
| DSF | 7/10/2019 | Diptera | Rhagionidae | * | 14430 (Bat156) | OL335054 | OL335112 | OL332646 | OL624654 |
| DSF | 8/1/2019 | Lepidoptera | Blastobasidae | * | 14431 (Bat160) | OL335055 | OL335113 | OL332647 | OL624655 |
| DSF | 8/1/2019 | Lepidoptera | Tineidae | *Dryadaula* sp. | 14448 (Bat163) | OL335056 | - | OL332648 | OL624656 |
| DSF | 8/1/2019 | Lepidoptera | Erebidae | *Lophocampa caryae* | 14435 (Bat164) | OL335057 | OL335114 | OL332649 | OL624657 |
| DSF | 8/1/2019 | Lepidoptera | Crambidae | *Eudonia* sp. | 14437 (Bat165) | OL335058 | OL335115 | OL332650 | OL624658 |
| DSF | 8/1/2019 | Lepidoptera | Blastobasidae | * | 14428 (Bat167) | OL335059 | - | OL332651 | OL624659 |
| DSF | 8/1/2019 | Diptera | Rhagionidae | * | 14432 (Bat173) | OL335060 | - | OL332652 | OL624660 |
| DSF | 8/1/2019 | Diptera | Dolichopodidae | *Thrypticus* sp. | 14436 (Bat174) | OL335061 | - | OL332653 | OL624661 |
| DSF | 8/1/2019 | Diptera | Sciaridae | * | (179) | OL335062 | OL335116 | OL332654 | OL624662 |
| DSF | 8/1/2019 | Diptera | Sciaridae | * | 14425 (Bat181) | OL335063 | OL335117 | OL332655 | OL624663 |
| DSF | 8/1/2019 | Diptera | Sciaridae | * | (Bat183) | OL335064 | OL335118 | OL332656 | OL624664 |
| DSF | 8/1/2019 | Diptera | Sciaridae | * | 14433 (Bat184) | OL335065 | OL335119 | OL332657 | OL624665 |
| DSF | 8/1/2019 | Diptera | Sciaridae | * | 14438 (Bat186) | OL335066 | OL335120 | OL332658 | OL624666 |
| DSF | 8/1/2019 | Diptera | Sciaridae | * | 14449 (Bat189) | OL335067 | OL335121 | OL332659 | - |
| DSF | 8/7/2019 | Coleoptera | Cantharidae | *Rhagonycha* sp. | 14434 (Bat204) | OL335068 | OL335122 | OL332660 | OL624667 |
| DSF | 8/7/2019 | Diptera | Sciaridae | * | 14439 (Bat205) | OL335069 | OL335123 | OL332661 | OL624668 |
| DSF | 8/7/2019 | Coleoptera | Cantharidae | *Rhagonycha* sp. | (Bat207) | OL335070 | OL335124 | OL332662 | OL624669 |
| DSF | 8/7/2019 | Hemiptera | Cixiidae | *Cixius* sp. | (Bat209) | OL335071 | OL335125 | OL332663 | OL624670 |
| DSF | 8/7/2019 | Lepidoptera | Blastobasidae | * | 14424 (Bat210) | OL335072 | - | OL332664 | OL624671 |
| DSF | 8/7/2019 | Coleoptera | Cantharidae | *Rhagonycha* sp. | (Bat217) | OL335073 | OL335126 | OL332665 | OL624672 |
| DSF | 8/7/2019 | Hemiptera | Derbidae | *Apache degeeri* | 14457 (Bat220) | OL335074 | OL335127 | OL332666 | OL624673 |
| DSF | 8/7/2019 | Lepidoptera | Oecophoridae | *Fabiola edithella* | 14440 (Bat221) | OL335075 | OL335128 | OL332667 | OL624674 |
| DSF | 8/15/2019 | Hemiptera | Cicadellidae | * | 14441 (Bat222) | OL335076 | OL335129 | OL332668 | OL624675 |
| DSF | 8/15/2019 | Lepidoptera | Erebidae | *Lymantria dispar* | (Bat228) | OL335077 | OL335130 | OL332669 | OL624676 |
| DSF | 8/15/2019 | Hemiptera | Cicadellidae | * | 14423 (Bat241) | OL335078 | OL335131 | OL332670 | OL624677 |
| DSF | 8/15/2019 | Hemiptera | Cixiidae | *Cixius* sp. | 14422 (Bat242) | - | OL335132 | OL332671 | OL624678 |
| DSF | 8/15/2019 | Diptera | Anthomyiidae | * | (Bat249) | OL335079 | OL335133 | OL332672 | OL624679 |
| DSF | 9/4/2019 | Diptera | Heleomyzidae | *Tephrochlamys rufiventris* | (Bat269) | OL335080 | OL335134 | OL332673 | OL624680 |
| DSF | 9/4/2019 | Psocoptera | Amphipsocidae | *Polypsocus corruptus* | (Bat270) | - | OL335135 | OL332674 | OL624681 |
| DSF | 9/4/2019 | Diptera | Sciaridae | * | 14450 (Bat271) | OL335081 | OL335136 | OL332675 | OL624682 |
| DSF | 9/4/2019 | Diptera | Psychodidae | * | 14451 (Bat273) | OL335082 | OL335137 | OL332676 | OL624683 |
| DSF | 9/4/2019 | Diptera | Dolichopodidae | *Gymnopterus* sp. | (Bat275) | - | - | OL332677 | - |
| DSF | 9/4/2019 | Hemiptera | Cixiidae | *Cixius* sp. | (Bat277) | - | - | - | OL624684 |
| DSF | 9/4/2019 | Hemiptera | Achilidae | * | (Bat287) | OL335083 | OL335138 | OL332678 | OL624685 |
| DSF | 9/4/2019 | Hemiptera | Achilidae | * | (Bat288) | OL335084 | OL335139 | OL332679 | OL624686 |
| DSF | 9/4/2019 | Hemiptera | Achilidae | * | (Bat291) | OL335085 | OL335140 | OL332680 | OL624687 |
| DSF | 9/14/2019 | Diptera | Sciaridae | * | (Bat297) | OL335086 | OL335141 | OL332681 | OL624688 |
| DSF | 9/14/2019 | Hemiptera | Cixiidae | *Cixius* sp. | 14458 (Bat300) | OL335087 | OL335142 | OL332682 | OL624689 |
| DSF | 9/14/2019 | Diptera | Sciaridae | * | (Bat301) | OL335088 | OL335143 | OL332683 | OL624690 |
| DSF | 9/14/2019 | Diptera | Sciaridae | * | 14452 (Bat304) | OL335089 | OL335144 | OL332684 | OL624691 |
| DSF | 9/14/2019 | Diptera | Sciaridae | * | 14453 (Bat309) | OL335090 | OL335145 | OL332685 | OL624692 |
| DSF | 9/14/2019 | Diptera | Sciaridae | * | 14429 (Bat320) | OL335091 | OL335146 | OL332686 | OL624693 |
| DSF | 9/14/2019 | Diptera | Sciaridae | * | 14454 (Bat321) | OL335092 | OL335147 | OL332687 | OL624694 |
| DSF | 9/14/2019 | Lepidoptera | Geometridae | *Lambdina fiscellaria* | (Bat322) | - | OL335148 | OL332688 | OL624695 |
| DSF | 9/14/2019 | Diptera | Sciaridae | * | 14455 (Bat323) | - | OL335149 | OL332689 | - |
| DSF | 9/14/2019 | Diptera | Sciaridae | * | (Bat324) | OL335093 | OL335150 | OL332690 | OL624696 |
| DSF | 9/14/2019 | Hemiptera | Derbidae | *Apache degeeri* | 14459 (Bat326) | OL335094 | OL335151 | OL332691 | OL624697 |
| AFF | 9/19/2019 | Diptera | Sciaridae | * | (Bat327) | OL335095 | OL335152 | OL332692 | OL624698 |
| AFF | 9/19/2019 | Diptera | Lauxaniidae | * | (Bat332) | OL335096 | OL335153 | OL332693 | - |
| AFF | 9/19/2019 | Diptera | Dolichopodidae | *Medetera* sp. | (Bat333) | - | OL335154 | OL332694 | OL624699 |
| AFF | 9/19/2019 | Diptera | * | * | (Bat334) | OL335097 | OL335155 | OL332695 | OL624700 |
| AFF | 9/19/2019 | Diptera | Milichiidae | *Madiza glabra* | (Bat335) | OL335098 | OL335156 | OL332696 | OL624701 |
| AFF | 9/19/2019 | Psocoptera | Psocomorpha | * | (Bat340) | OL335099 | OL335157 | OL332697 | OL624702 |
| AFF | 8/31/2020 | Hemiptera | Fulgoridae | *Lycorma delicatula* | (Bat565) | OL335100 | OL335158 | OL332698 | OL624703 |

AFF - Angora Fruit Farm, Berks County Parks and Recreation, Pennsylvania, DSF – Danby State Forest, Tompkins County, New York. – missing data, \* - not identified to the family/suborder, genus, or species level.

### DNA extraction and amplification

Fungal tissues from in vitro growth were transferred to lysis buffer and beaten with 0.5 g of 0.7 mm diameter zirconia beads at 4800 rpm for 1 min. DNA extraction and PCR were performed as described in Hajek et al. (15). PCR was performed on 4 loci: 28S, ITS1, ITS2 and *RPB2*. 28S amplification used forward primer LR0R (16) and reverse primer LR5 (17). ITS1 amplification used forward primer ITS5 (18) and reverse primer 5.8S (17). ITS2 amplification used forward primer ITS3 (18) and reverse primer ITS4sub: 5’-TGGAGCAAGTACAAACAACACT-3’. *RPB2* amplification used forward primer BatRPB2f: 5’-ACCCTCAGAAACCTCTCGTC-3’ and reverse primer BatRPB2r: 5’-CAAACCGAGCCAGCAATTTG-3’.

PCR conditions for 28S were initial denaturation for 5 min at 95°C followed by 6 cycles of denaturation for 1 min at 95°C, annealing at 58°C for 1 min that decreased by 1°C for each cycle, and extension for 1.5 minutes at 72°C. The 6 cycles were followed by 30 cycles of denaturation for 30 sec at 95°C, annealing at 52°C for 1 min, and extension for 1 min at 72°C. The final step was extension at 72°C for 10 sec. PCR conditions for ITSI and ITSII were an initial denaturation for 5 min at 94°C followed by 35 cycles of denaturation at 94°C for 45 sec, annealing at 55°C for 50 sec, and extension at 72°C for 1 min. The final step was extension at 72°C for 10 min. PCR conditions for *RPB2* were an initial denaturation at 95°C for 4 min followed by 34 cycles of denaturation at 95°C for 1 min, annealing at 50°C for 1 min, a ramp that increased the temperature at a rate of 0.3°C/sec for 1.23 min from 50 to 72°C, and extension for 1 min at 72°C. The final step was extension for 10 min at 72°C (19). To check whether the PCR products were viable, products underwent agarose gel electrophoresis in 1x TAE buffer and were visualized with ACCURIS SmartDoc (Accuris, New Jersey, USA). Successful products were purified by combining 8μL of product with 2μL of a master mix (1.6μL molecular water, 0.2μL 10X PCR buffer, 0.1μL SAP enzyme, and 0.1μL EXO enzyme) at 37°C for 35 min followed by deactivating the enzymes at 90°C for 13 min. Purified PCR products were sequenced by Genewiz LLC (South Plainfield, New Jersey, USA). Sequences were edited, assembled, aligned, and searched using Geneious software v. 8.1.8 (Biomatters Ltd).

### Phylogenetic reconstruction and analysis of population structure

A single FASTA file was prepared from each of the four loci used to identify *B. major*: ITS1(N=39), ITS2 (N=54), 28S (N=66), and *RPB2* (N=62). Each FASTA file was aligned using MAFFT version 7 (using default parameters with a scoring matrix of 1PAM/ κ=2 for closely related sequences) and was imported into R version 4.0.2 (20). All analyses were conducted using the *adegenet* and *poppr* packages (21,22). Fungal samples in each FASTA file were further labelled according to the geographic location and arthropod host from which they were collected (accounting for host order, family, and species).

To infer the number of genetic clusters across our data set, and to evaluate the utility of arthropod host as a predictor of population structure in *B. major*, alignments of each locus were subjected to a Discriminant Analysis of Principle Components (DAPC) (23–25). An additional DAPC was performed retaining only one sample per genotype per population using the clonecorrect command in poppr (22). Population differentiation was further analyzed by calculating FST according to *B. major* host order and family using hierfstat (26).

## Results

### Morphological characterization

Our morphological observations of *B. major* from the cadavers of host insects do not differ from previously published records (2). Diameters of conidia, size of conidial papillae, and the number of nuclei in conidia are typical for the species (Fig 1).

**Fig 1.**
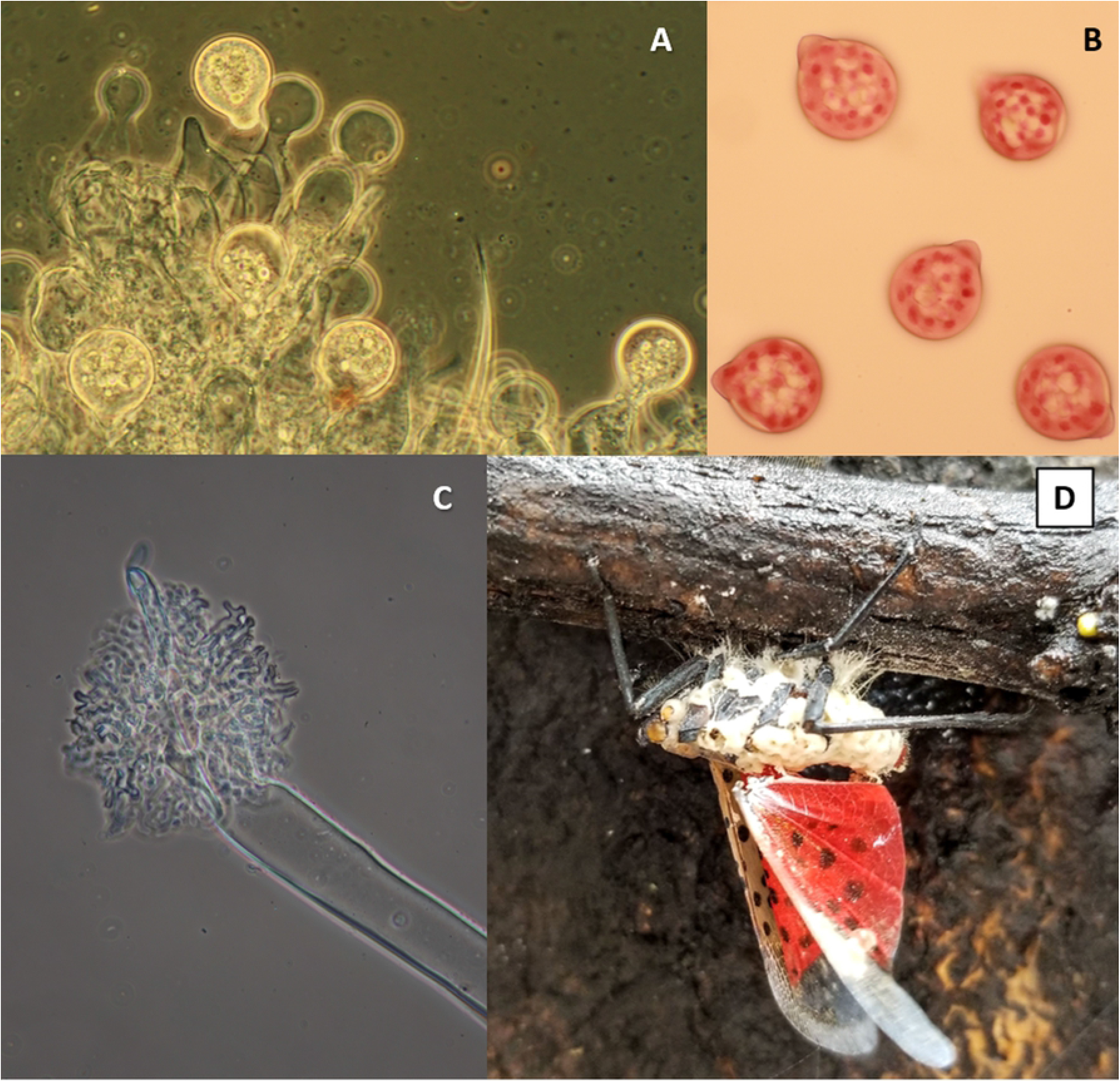
Micromorphology of *Batkoa major*. **A**. Conidiophores with conidia. Conidia average 41.5 µm wide x 49.3 µm long. **B**. Multinucleate conidia (nuclei stained with aceto-orcein). **C**. Distal end of rhizoid with holdfast. **D**. Cadaver of the spotted lanternfly attached to a twig by rhizoids (Photo by E.H. Clifton).

### Genetic polymorphism reveals the lack of host specificity

Genotype comparisons and phylogenetic reconstructions suggests that these *B. major* populations consist of numerous genotypes. Alignments of four loci (28S, ITS1, ITS2 and *RPB2*) of *B. major* reveal a high degree of genetic polymorphism among individuals (Fig 2).

**Fig 2.**
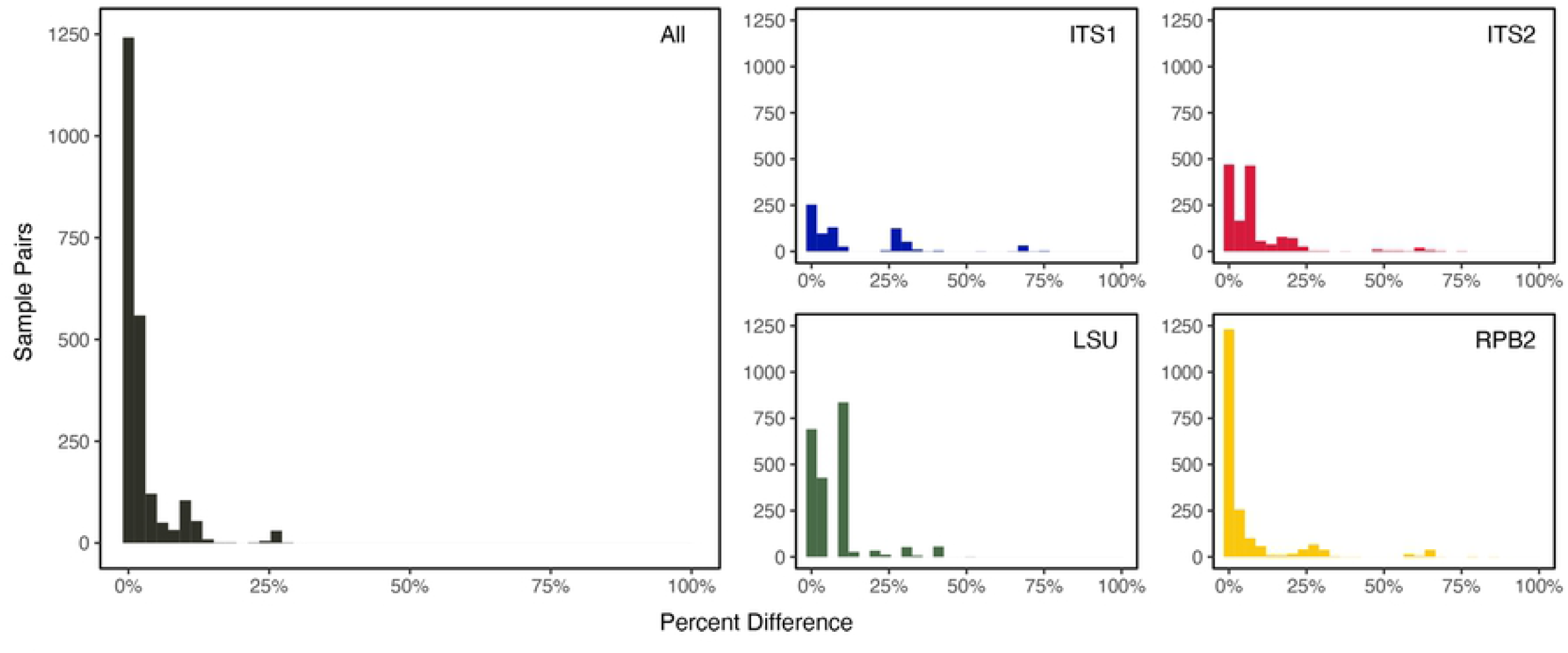
Percent genetic difference among pairwise comparisons of all specimens.

Approximately half of all specimen pairs are identical for any given locus, as well as when all four loci are combined, suggesting that asexuality is an important aspect of the *B. major* life history. The least degree of polymorphism occurs in *RPB2*, whereas the most occurs in ITS1, in contrast to observations in other fungal species (27,28).

The combined alignments consisted of 3,201 characters (N=67 specimens), with each locus presenting a varied degree of sites: 28S had a length of 1,067 bp with 20 polymorphic positions (N=66), ITS1 had a length of 911 bp with 174 polymorphic positions (N=39), ITS2 had a length of 639 bp with 58 polymorphic positions (N=54), and *RPB2* had a length of 584 bp with 29 polymorphic positions (N=62). Although we were unable to amplify and sequence the entire ITS region, and thus combine ITS1 and ITS2 sequences, we estimate the total length of the ITS region is greater than 1,600 characters.

Notably, genetic polymorphism does not correspond to arthropod host (Fig 3). For example, several clonal sample (i.e., those with branch lengths of zero in Fig 3) were collected from arthropod hosts distantly related to each other and belonging to different orders. Even when genetically dissimilar specimens are compared, several host orders or families are represented in a single clade.

**Fig 3.**
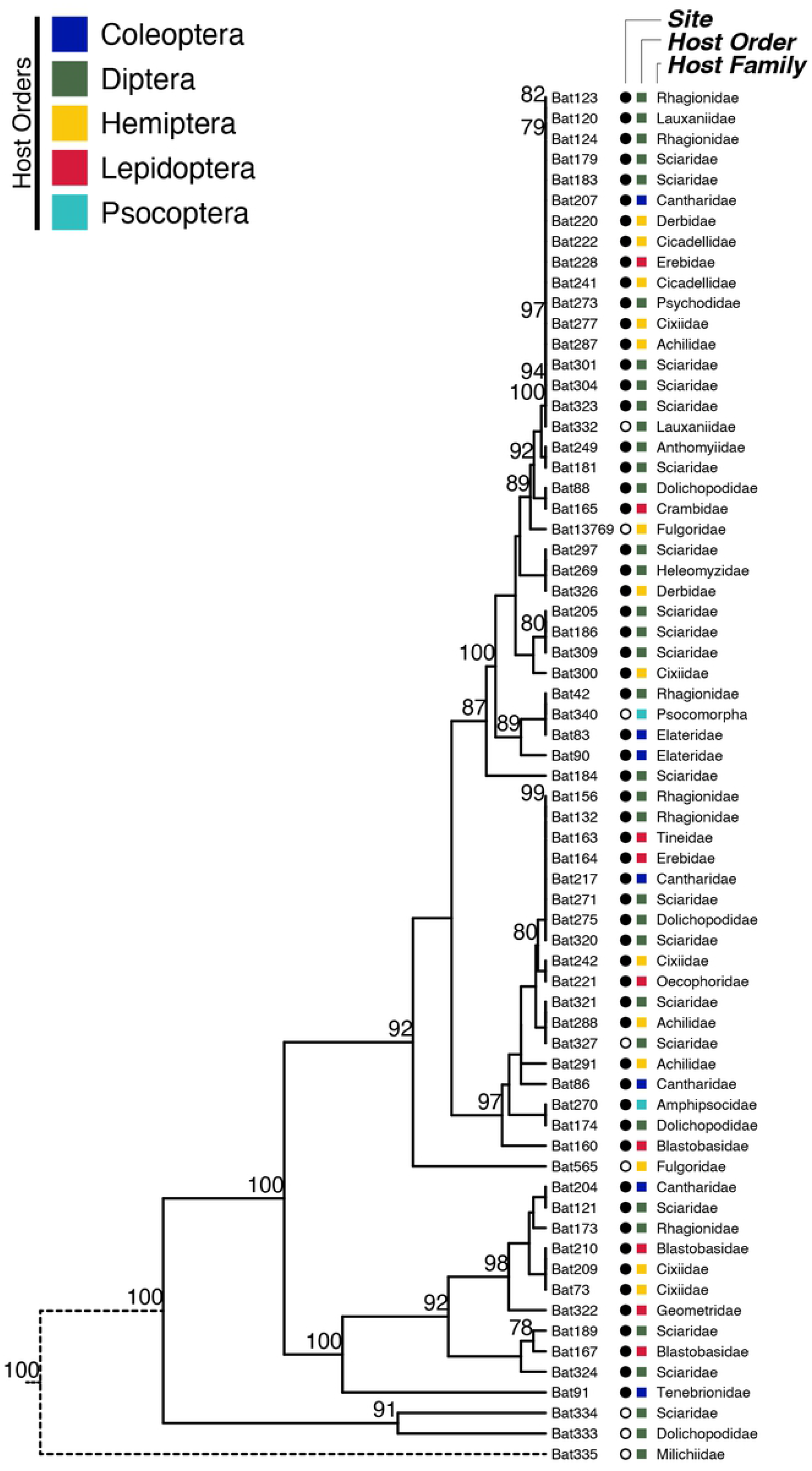
Dendrogram of all *B. major* isolates. Each specimen is labelled according to the order and family of the arthropod host from which it was sampled. Population origin is also specified, with an open circle for the Angora Fruit Farm population and a closed circle for the Danby State Forest population. Nodes whose bootstrap support was greater than 75 are labelled accordingly. A dotted black line was drawn to depict the outermost tree branch to indicate that it was shortened for aesthetic purposes.

Mapping hosts on the *B. major* phylogenetic tree demonstrated that hosts from the same family can be infected by pathogens that have different genotypes. There is no visible grouping of the hosts with particular clades of the pathogen, either on the single locus trees (Suppl Fig 1) or on the combined four-locus tree (Fig 3). Major host orders and families are located on the *B. major* phylogenetic tree randomly. Slightly more than half of the infected insects belong to order Diptera (35 out of 67 samples). Among individual families, the most samples were from the Sciaridae (dark-winged fungus gnats; 19 out of 67 samples). Insect orders attacked less frequently by *B. major* are (in descending order): Hemiptera (N=14 samples), Lepidoptera (N=9), Coleoptera (N=7), Psocoptera (N=2).

Our DAPC analysis corroborates a lack of host specificity in *B. major*, illustrating a high degree of genotypic overlap when grouping samples by host order. The first (44.2% [64.1% clone corrected]) and second (26.2% [22.5% clone corrected]) principal components capture most of the genetic variation among samples and reveal no apparent clustering of individuals in correlation to their arthropod host (including when only a single specimen per genotype is retained in the analysis). An exception was the two samples from the Order Psocoptera that clustered together in the clone-corrected analysis (Fig 4).

**Fig 4.**
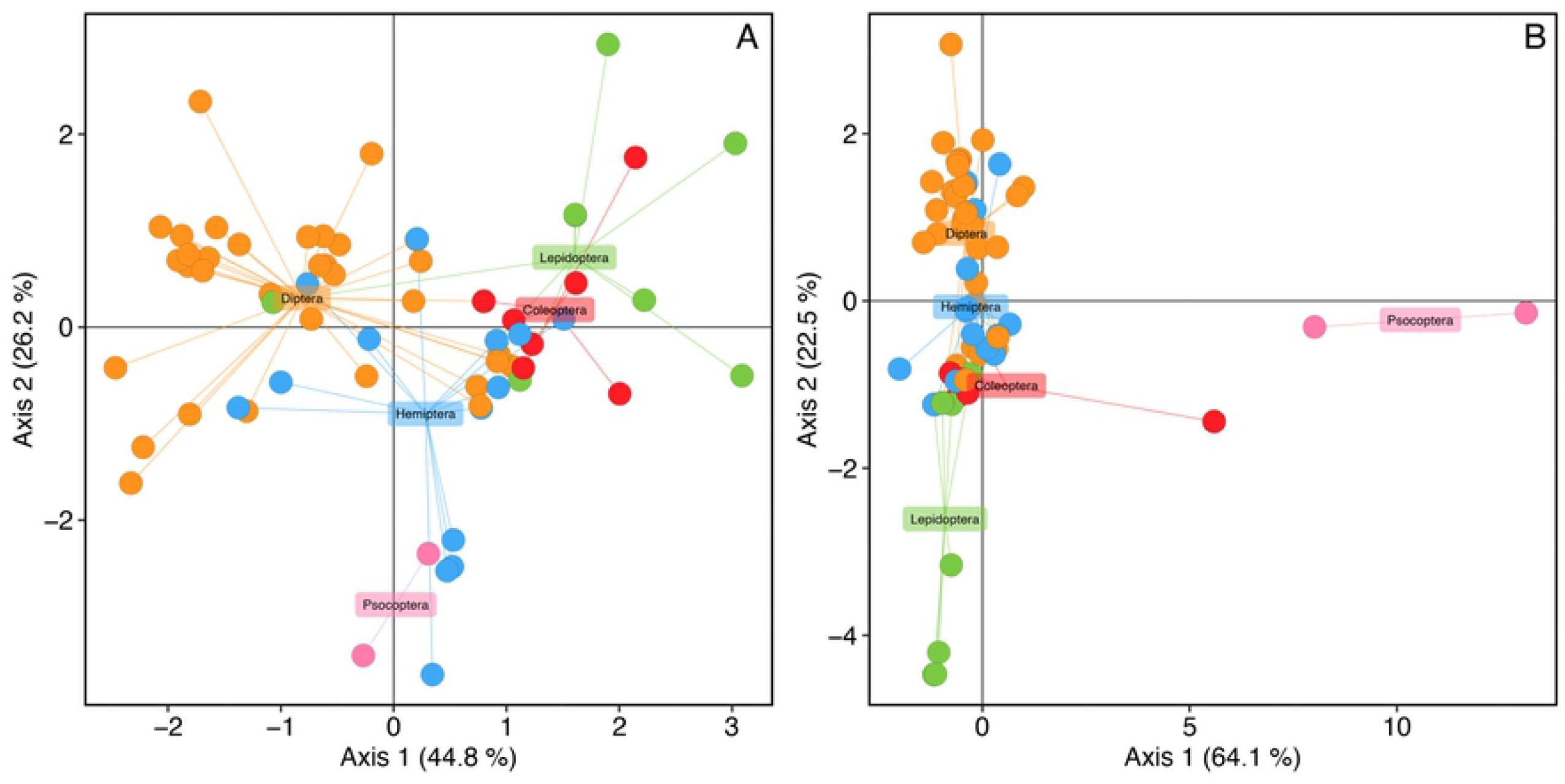
DAPC analysis for the combined ITS1, ITS2, LSU and *RPB2* Data in full (**A**) and clone-corrected (**B**). Specimens are labeled according to host order. Axis 1 explained 44.8% (64.1% for clone-corrected data) and axis 2 explained 26.2% (22.5% for clone-corrected data) of the genetic variation among individuals.

### High gene flow within and between populations of *B. major*

Genotypes of *B. major* identified with either ITS1, ITS2, 28S or *RPB2* do not cluster according to collection site, in addition to not clustering by host order or family (Figs 3 and 4). Despite differences in sample size from our two populations, hosts were infected (and often belonging to different arthropod orders and families) at each location by the same genotype, despite being separated by 220 km. This suggests that dispersal via asexual spores or hyphal fragments contributes to the long-distance movement of *B. major* genotypes. Moreover, genetically distinct fungal specimens from the same population do not cluster phylogenetically, illustrating a broader pattern of high gene flow among genotypes of *B. major*.

The lack of population structure by arthropod host is further supported by FST values calculated between host order and family, in addition to values calculated between the two collection sites (Fig 5). In all cases, median F_ST_ is 0.05-0.035 in all pairwise comparisons of host orders and families, with the maximum F_ST_ values approximately 0.20 when comparing host family by the 28S locus.

**Fig 5.**
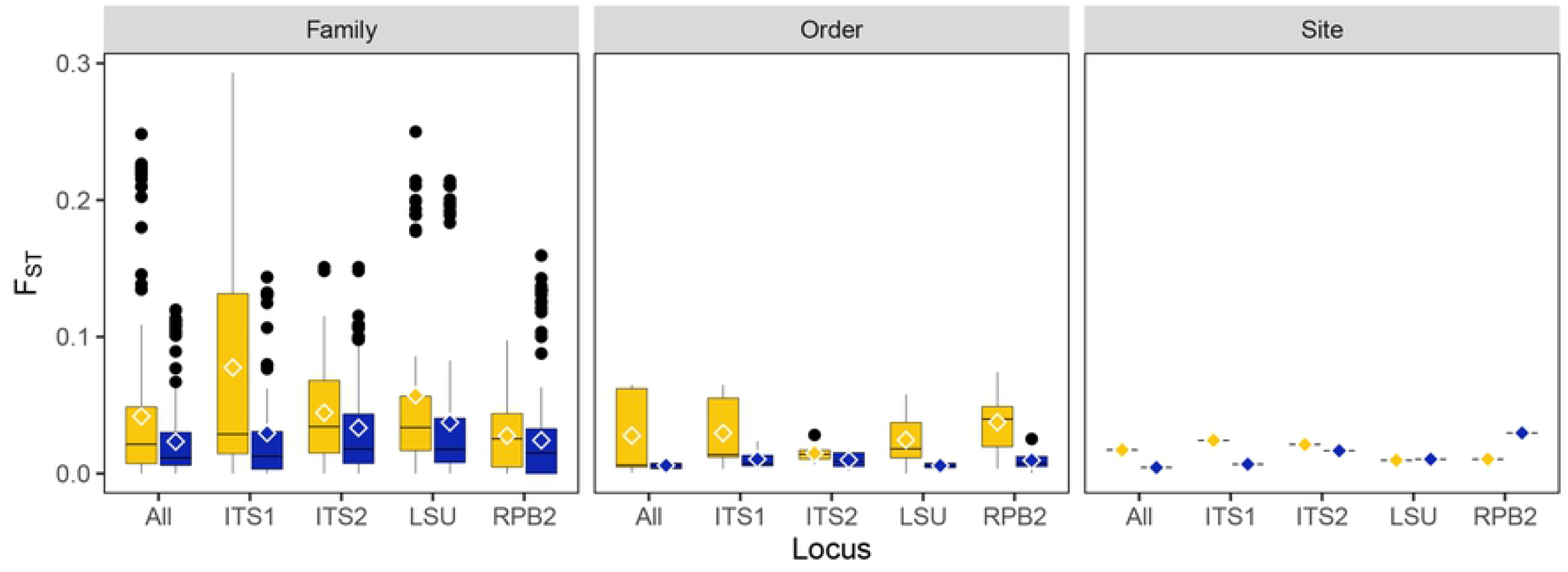
F_ST_ calculations for each locus and all four combined. F_ST_ calculated between host order, host family, and collection site (syn. population). Yellow – all data, blue – clone corrected data. Mean F_ST_ is illustrated by an open rhombus.

Interestingly, F_ST_ values indicate that not only is there high gene flow among populations, but also that there is high gene flow within populations vis-à-vis arthropod host. More specifically, our data not only indicate a lack of host specificity, but also that genotypes of *B. major* readily exchange genetic information (i.e., undergo sexual reproduction) with other genotypes infecting a phylogenetically distant arthropod host.

## Discussion

Entomophthoralean species have several different modes of host range diversity. *Batkoa major, B. apiculata, Zoophthora radicans*, and *Conidiobolus thromboides* have broader host ranges that include hosts in different insect orders. Then, there are some species of Entomophthorales with an intermediate type of host specificity, only infecting insects within the same insect order. For example, studies of physiological host range demonstrated that all three species in the *Entomophaga aulicae* species complex infect only species of Lepidoptera (29,30). Finally, there are highly specialized species, like *Strongwellsea magna* and *S. castrans*, that, even with extensive study, are known only from host species within individual families of Diptera (31). The group of species with broad host ranges seems to be the smallest group within this fungal order (2). Theory has suggested that more host specific parasites can lead to greater survival success (32).

Contrary to expectations, we found that the native fungal entomopathogen *B. major* has a diverse host range, including native insects in five insect orders. Why this fungal species has not been reported more previously is not known although one possibility is the difficulty of sampling insects in forested locations. Nonetheless, our findings agree with those for other fungal pathogens with broad host ranges, which are generally held to be more likely to form symbioses with novel hosts in invasive contexts (33,34).

Invasive *L. delicatula* is thus a competent host for this native pathogen that is a generalist and which we found in abundance during a fall epizootic in *L. delicatula* (13) or all season long in different native host species. The well-known idiom ‘jack of all trades and master of none’ has been used to suggest that generalists would be less successful than specialists (32). The alternate opinion that changes the idiom to ‘jack of all trades and master of all’ (35) is more consistent with results from the present study where *B. major* was found all season long infecting a diversity of hosts, although prevalence was not high in these lower density populations. Woolhouse et al. (36) suggest that conditions predisposing pathogens to generalism include high levels of genetic diversity as well as ample opportunities for cross-species transmission. While we found clonality for some groups of *B. major*, we found numerous clones including multiple samples from different hosts and gene flow occurred among some of them. We see some similarity in the population structure between *B. major* (our observations) and *E. muscae* (37). In addition, our predominant collecting site was a native forest in New York during summer and the native insect fauna provided a diversity of hosts. Clones also existed within each of two clades of the entomophthoralean *Entomophthora muscae* infecting two species of flies (38).

Unusually rDNA sequence length significantly contributes to the polymorphism in our samples, possibly also due to higher substitution rates compared to other entomophthoralean fungi (9). Curiously, the total length of the ITS region in *B. major* exceeds 1600 bp, which is quite an unusual feature compared to most fungal species. However, long ITS is also characteristic of other entomophthoralean species, e.g. for *Entomophthora muscae* (39) and *Zoophthora* species (40). Also, genomes in some fungi contain multiple ITS copies (41). Therefore, high population diversity in the *B. major* population recorded for the ITS1 and ITS2 regions might significantly reflect random ITS copies rather than real genetic diversity. The longer that the length of the ITS region is, the larger the number of mutations that might occur and therefore the larger number of different ITS copies that might be amplified and sequenced, which can be reflected as population diversity for that genomic region. In contrast, high numbers of identical copies suggest a high degree of clonality in the population, i.e., identical copies with different placement on the phylogenetic tree. It seems unlikely that we have randomly sampled the same ITS copy. This fact might be a good indication that the copies in *B. major* are homogenized by concerted evolution and sexual processes are occurring.

Generalist pathogens are thought to potentially experience trade-offs in that they are not as well adapted to all the hosts that they infect (36). While Bufford et al. (34) found that taxonomic similarity of co-evolved hosts with novel hosts was more important than contact opportunity, our study did not find any such patterns. In the present study, it could be possible that fitness could differ when *B. major* infects the invasive *L. delicatula* versus the diverse native hosts that were infected. Similar observations were made for the efficiency of *E. muscae* infecting even closely related muscoid species at the same location (42). As opposed to the present study, the clones in *E. muscae* were associated with host species. For the fungal clavicipitacean genus *Metarhizium*, clones occurred within different species; however, because isolates came from soil samples, host relationships are not possible (43).

Yet, even if individual fitness was decreased when *B. major* infected the novel invasive *L. delicatula*, being a generalist allowed *B. major* to take advantage of an outbreak population of an invasive host and we did not find native specialist pathogens responding to these outbreak invasive populations.

## Conclusion

The studied populations of *B. major* can infect various hosts in the same location. Analysis of molecular data supports the hypothesis of the clonal nature of the studied population. This can serve as a good example of a genetically diverse population of a pathogen-generalist with a certain amount of gene flow between its members. Use of a broad host range enabled *B. major* to switch to infection of the spotted lanternfly, a new invasive pest in the USA, which only appeared in Pennsylvania in 2014.

## Acknowledgments

We thank David Harris and Sarah Stefanik for collection, isolation, and identification of fungi. We thank Jim Liebherr, Tyler Hagerty, Charles Bartlett, Jason Dombroskie for identification of some fungal-killed insects. This research was supported by USDA NIFA SCRI 2019-51181-30014.

## Supporting Information

**S1 Fig**. Absence of visible grouping of the hosts with particular clades of the pathogen on the single locus trees for ITS1 (S1 A), ITS2 (S1 B), 28S (S1 C), and RPB2 trees (S1 D).

